# A comparison of blood and brain-derived ageing and inflammation-related DNA methylation signatures and their association with microglial burdens

**DOI:** 10.1101/2020.11.30.404228

**Authors:** Anna J. Stevenson, Daniel L. McCartney, Gemma L. Shireby, Robert F. Hillary, Declan King, Makis Tzioras, Nicola Wrobel, Sarah McCafferty, Lee Murphy, Barry W. McColl, Paul Redmond, Adele M. Taylor, Sarah E. Harris, Tom C. Russ, Eilis J Hannon, Andrew M. McIntosh, Jonathan Mill, Colin Smith, Ian J. Deary, Simon R. Cox, Riccardo E. Marioni, Tara L. Spires-Jones

## Abstract

Inflammation and ageing-related DNA methylation patterns in the blood have been linked to a variety of morbidities, including cognitive decline and neurodegenerative disease. However, it is unclear how these blood-based patterns relate to patterns within the brain, and how each associates with central cellular profiles. In this study, we profiled DNA methylation in both the blood and in five *post-mortem* brain regions (BA17, BA20/21, BA24, BA46 and hippocampus) in 14 individuals from the Lothian Birth Cohort 1936. Microglial burdens were additionally quantified in the same brain regions. DNA methylation signatures of five epigenetic ageing biomarkers (‘epigenetic clocks’), and two inflammatory biomarkers (DNA methylation proxies for C-reactive protein and interleukin-6) were compared across tissues and regions. Divergent correlations between the inflammation and ageing signatures in the blood and brain were identified, depending on region assessed. Four out of the five assessed epigenetic age acceleration measures were found to be highest in the hippocampus (β range=0.83-1.14, p≤0.02). The inflammation-related DNA methylation signatures showed no clear variation across brain regions. Reactive microglial burdens were found to be highest in the hippocampus (β=1.32, p=5×10^-4^); however, the only association identified between the blood- and brain-based methylation signatures and microglia was a significant positive association with acceleration of one epigenetic clock (termed DNAm PhenoAge) averaged over all five brain regions (β=0.40, p=0.002). This work highlights a potential vulnerability of the hippocampus to epigenetic ageing and provides preliminary evidence of a relationship between DNA methylation signatures in the brain and differences in microglial burdens.

## 1. Introduction

Ageing is characterised by a progressive deterioration of physiological integrity and is a key risk factor for a multitude of diseases. A pervasive feature of ageing is a persistent, or chronic, systemic inflammation (1). This process is characterised by a subtle elevation of inflammatory mediators in the periphery, in the absence of evident precipitants or disease states. Chronic inflammation has been identified as a common feature in the preponderance of neurodegenerative diseases and is increasingly recognised as a potential mediator of cognitive impairment in older age (2). There is, however, still a paucity of understanding of the biological mechanisms involved in chronic inflammation and how peripheral and central inflammatory mechanisms relate.

Recently, the link between inflammation and the epigenetic mechanism of DNA methylation (DNAm) has begun to be addressed (3, 4). DNAm is typically characterised by the addition of a methyl group to a cytosine, in the context of a cytosine-guanine (CpG) dinucleotide. It has been implicated in the regulation of gene expression and can itself be influenced by both genetic and environmental factors (5, 6). Genome-wide DNAm patterns in the blood have been leveraged to index lifestyle traits, such as smoking (7, 8), and have been used to investigate diverse physical and mental health-related phenotypes, including cognitive functioning (9). In addition to this, by exploiting the manifest alterations in DNAm patterns with ageing, several DNAm-based markers of age have been developed, which attempt to provide surrogate measures of biological ageing (10–13). These ‘epigenetic clocks’ have been used to provide a measure of biological age acceleration, or deceleration, by establishing the difference between an individual’s chronological and epigenetic age. Positive age acceleration quantified in the blood has been associated with an increased risk of mortality and a variety of age-related morbidities, including with a lower cognitive ability (14–16). In addition to this, recently, we found that blood-based DNAm proxies for two inflammatory mediators – C-reactive protein (CRP) and interleukin-6 (IL-6) – were inversely associated with cognitive ability in older adults with larger effect sizes compared to the biomarkers themselves (17, 18).

While these findings suggest that an accelerated biological age, and raised DNAm inflammation patterns associate with poorer cognitive functioning, it is important to note that these studies analysed blood tissue. While the blood represents a practical, accessible source by which to investigate such outcomes, DNAm is known to confer both cell-type and tissue-specific patterns (19). For analyses of brain-based traits such as cognitive ability, brain samples offer the optimal diseaserelevant tissue; however, given the obvious limitations of access to such tissue, much of the research assessing the association between differential DNAm and disorders of the central nervous system has been conducted in peripheral whole blood (20, 21). While this approach can provide informative peripheral markers of central aberration or disease, it is important to investigate the relevant target tissue to characterise both how peripheral and central patterns equate, and how each relates to cellular differences within the brain. Microglia are the primary tissue-resident immune cells of the central nervous system and have critical roles in homeostasis and neuroinflammation. Aged microglia have been shown to be more responsive to pro-inflammatory stimuli compared to naïve microglia, and evidence suggests the cells are particularly sensitive to both acute and chronic systemic inflammation detected via peripheral-central signalling pathways (22, 23). Microglia have additionally been implicated in age-related neurological dysfunction; however, as yet, it is unclear how inflammation and age-related DNAm patterns in both the periphery and the brain itself relate to microglial burdens.

In this study, we utilise data from 14 participants of the Lothian Birth Cohort 1936. These individuals have blood-based DNAm data available at up to 4 time-points between the ages of 70-79 years and additionally donated *post-mortem* brain tissue to the study. In the brain, we profiled DNAm and quantified microglial burdens in five regions (primary visual cortex [BA17], inferior temporal gyrus [BA20/21], anterior cingulate cortex [BA24], dorsolateral prefrontal cortex [BA46], and hippocampus). DNAm CRP and IL-6 profiles, along with five different DNAm age acceleration measures, were characterised in the blood and in each brain region to investigate the relationship between peripheral and central age- and inflammation-related methylation patterns and how these relate to inflammatory processes in the brain. Given the small sample size of this study, the results presented here represent preliminary patterns; however, this data, and the methodology employed, provides a framework upon which future, larger scale, work can be based.

## 2. Methods

### 2.1 The Lothian Birth Cohort 1936

The Lothian Birth Cohort 1936 (LBC1936) is a longitudinal study of ageing. Full details on the study protocol and data collection have been described previously (24, 25). Briefly, the cohort comprises 1,091 individuals born in 1936 most of whom completed a study of general intelligence – the Scottish Mental Survey – in 1947 when they were aged around 11 years. Participants who were living in Edinburgh and the surrounding area were re-contacted around 60 years later with 1,091 individuals consenting to join the LBC1936 study. At Wave 1 of the study participants were around 70 years old (mean age: 69.6□±□0.8 years) and they have since completed up to four additional assessments, triennially. At each assessment, participants have been widely phenotyped with detailed physical, cognitive, epigenetic, health and lifestyle data collected. A tissue bank for *post-mortem* brain tissue donation was established at Wave 3 of LBC1936 in collaboration with the Medical Research Council-funded University of Edinburgh Brain Banks. To date, ~15% of the original LBC1936 sample have given consent for *post-mortem* tissue collection. At the time of this study, samples from 14 individuals were available.

### 2.2 Ethics

Ethical permission for LBC1936 was obtained from the Multi-Centre Research Ethics Committee for Scotland (MREC/01/0/56), the Lothian Research Ethics Committee (Wave 1: LREC/2003/2/29) and the Scotland A Research Ethics Committee (Waves 2, 3 and 4: 07/MRE00/58).

Use of human tissue for *post-mortem* studies was reviewed and approved by the Edinburgh Brain Bank ethics committee and the medical research ethics committee (the Academic and Clinical Central Office for Research and Development, a joint office of the University of Edinburgh and NHS Lothian, approval number 15-HV-016). The Edinburgh Brain Bank is a Medical Research Council funded facility with research ethics committee (REC) approval (16/ES/0084).

### 2.3 DNA methylation preparation

#### 2.3.1 Blood

DNAm from whole blood was quantified at 485,512 CpG sites using the Illumina Human Methylation 450k BeadChips at the Edinburgh Clinical Research Facility. Full details of the quality control steps have been described previously (26, 27). Briefly, raw intensity data were background-corrected and normalised using internal controls. Samples with inadequate bisulphite conversion, hybridisation, staining signal or nucleotide extension were removed upon manual inspection. Further, probes with a low detection rate (p>0.01 in >5% of samples), samples with a low call rate (<450,000 probes detected at p<0.01), samples exhibiting a poor match between genotype and SNP control probes, and samples with a mismatch between methylation-predicted, and recorded, sex were additionally excluded. This left a total of 450,276 autosomal probes. In analyses comparing blood and brain DNAm signatures, the last blood measurement before death was used and models were adjusted for the interval between the blood draw and death (see **Supplementary Table** 1; mean interval: 2.5 years, SD: 1.5).

#### 2.3.2 Brain

Brains were removed at *post-mortem* and cut into coronal slices. Regions of interest dissected, as detailed previously (28). Tissue samples from cortical regions BA17, BA20-21, BA24, BA46 and hippocampus, were collected and snap frozen. From these sections ~25mg of tissue was processed for DNA extraction. DNA extraction was performed using a DNeasy kit (Qiagen) and DNAm was profiled using Illumina MethylationEPIC BeadChips at the Edinburgh Clinical Research Facility. Samples were processed randomly. Quality control steps were performed as follows: the *wateRmelon* pfilter() function (29) was used to remove samples in which >1% of probes had a detection p-value of >0.05, probes with a beadcount of <3 in >5% of samples, and probes in which >1% of samples had a detection p-value of >0.05. Probes mapping to polymorphic targets, cross-hybridising probes and probes on the X and Y chromosomes were additionally removed. The performance of 15 normalisation functions was assessed, following the protocol described by Pidsley *et αl.* (29). The top-ranking method was *danet* which equalises background from type 1 and type 2 probes, performs quantile normalisation of methylated and un-methylated intensities simultaneously, and then calculates normalised methylation β-values. The normalised dataset comprised 69 samples (14 individuals, 5 regions, 1 missing hippocampal sample) and 807,163 probes.

### 2.4 Derivation of DNA methylation signatures

#### 2.4.1 Epigenetic age acceleration

Methylation-based epigenetic age acceleration estimates were obtained from the online Horvath DNAm age calculator (https://dnamage.genetics.ucla.edu/)(11). Normalised DNAm data was uploaded to the calculator using the ‘Advanced Analysis’ option. This output provides four different age acceleration measures: intrinsic epigenetic age acceleration (IEAA) (11); extrinsic epigenetic age acceleration (EEAA) (12); DNAm PhenoAge acceleration (AgeAccel_pheno_)(10); and DNAm GrimAge acceleration (AgeAccel_Grim_)(13). IEAA is defined as the residuals resulting from the regression of estimated epigenetic age based on the Horvath epigenetic clock on chronological age, fitting estimated proportions of immune cells. IEAA is designed to capture cell-intrinsic epigenetic ageing, independent of age-related changes in blood cellular composition. EEAA is estimated firstly by calculating a weighted average of Hannum’s methylation age with three cell types — naïve cytotoxic T cells, exhausted cytotoxic T cells and plasmablasts. EEAA is defined as the residuals resulting from the univariate regression of this weighted estimate on chronological age and correlates with age-related changes in the blood cellular composition. Though these measures are most appropriate for use in the blood as they account for blood cell proportions, the correlation between these and the unadjusted measures are both >0.97, suggesting they are very similar. Rather than aiming to predict chronological age, DNAm PhenoAge was designed to capture an individual’s ‘phenotypic age’ – a composite set of clinical measures associated with mortality. Regressing DNAm PhenoAge onto chronological age provides the acceleration measure: AgeAccel_pheno_. Similarly, DNAm GrimAge was designed to predict mortality based on a linear combination of age, sex, and DNAm-based surrogates for smoking and seven plasma proteins. AgeAccel_Grim_ provides the measure of epigenetic age acceleration from this clock. In addition to the epigenetic age acceleration measures, the online calculator provides an estimate of the proportion of neurons in each sample, derived using the cell epigenotype specific (CETS) algorithm (30).

Recently, a novel epigenetic clock (DNAmClock_Cortical_) was developed to optimally capture brain-specific epigenetic ageing (31). This clock was trained on 9 human cortex methylation datasets of tissue from individuals unaffected by Alzheimer’s disease (total n=1,397, age range=1-104 years). The model selected 347 DNAm sites and the clock was then tested in an external cohort, outperforming other epigenetic clocks for age prediction within the brain. The sum of DNAm levels at these sites weighted by their regression coefficients provided the cortical DNAmClock_Cortical_ age estimate. The residuals resulting from regressing DNAmClock_Cortical_ age on chronological age provided the age acceleration measure for this epigenetic clock (AgeAccel_Cortical_).

#### 2.4.2 Inflammation signatures

DNAm scores for the acute-phase inflammatory mediator C-reactive protein (CRP) and the pro-inflammatory cytokine interleukin-6 (IL-6) were derived as described previously (17, 18, 32). The DNAm CRP score was obtained using data from a large epigenome-wide association study (EWAS) of CRP (3). This EWAS identified 7 CpG sites with strong evidence of a functional association with circulating CRP. One of these CpGs (cg06126421) was not available on the EPIC array, therefore the sum of DNAm levels at the remaining 6 CpG sites weighted by their regression coefficients from the EWAS provided the DNAm CRP score (32) **(Supplementary Table 2).** The IL-6 score was derived from an elastic net penalised regression model using the Wave 1 LBC1936 blood methylation and Olink^®^ IL-6 data (Olink^®^ inflammation panel, Olink^®^ Bioscience, Uppsala, Sweden) (17). This approach identified 12 CpG sites that optimally predicted circulating IL-6. In the current study, the elastic net regression was re-run omitting individuals providing *post-mortem* brain samples (n=863). This model returned a set of 13 CpG sites (inclusive of the 12 CpGs from the original model). The DNAm IL-6 score in both blood and brain were thus derived from the sum of DNAm levels at these 13 CpG sites weighted by their regression coefficients **(Supplementary Table 3).**

### 2.5 Immunohistochemistry, thresholding and burden quantification

Fixed tissue sections (4μm) from cortical regions BA17, BA20-21, BA24, BA46 and hippocampus were processed for immunohistochemistry. Staining was carried out as described previously (33). Briefly, microglial lysosomes were stained using CD68 (mouse anti-human monoclonal primary antibody, Dako M0876,1:100, citric acid in pressure cooker pre-treatment). Immunohistochemistry was performed using standard protocols, enhanced with the Novolink Polymer detection system and visualised using 3,3’-diaminobenzidine (DAB) with 0.05% hydrogen peroxide as chromogen. Tissue was counterstained with haematoxylin for 30 seconds to visualise cell nuclei.

Stains were visualised using a ZEISS Imager.Z2 stereology microscope using MBF Biosciences Stereo Investigator software. All 6 layers of cortical grey matter were included in analysis. Cortical grey matter was outlined at 1.5X objective magnification and tile scans were acquired at 5X for quantification. Glia were quantified using in-built software that captures immuno-positive objects using an automated thresholding algorithm based on colour and size. Objects smaller than 10μm^2^ were not considered true staining and were thus excluded in the burden analysis. The threshold and exposure remained consistent throughout all analysis. Neurolucida Explorer was used to quantify the total area of the region of interest and that of the outlined objects. A percentage burden was then calculated by dividing the stained area by the total tissue area.

### 2.6 Statistical analyses

Spearman correlations were calculated between the inflammation, and epigenetic age acceleration, measures in the blood and each brain region using the last available blood-based measure prior to death. Linear mixed effects models were used to investigate the regional heterogeneity in the epigenetic age acceleration variables and the DNAm inflammation scores in the brain. BA17 was set as the reference as this region is typically not affected until the latter stages of neurodegenerative diseases that impact cognitive functioning, such as Alzheimer’s disease. Models were adjusted for age at death, *post-mortem* interval, sex, and proportion of neurons, with participant ID fitted as a random effect on the intercept. Linear mixed effects models were additionally used to assess the association between the DNAm signatures in both the blood and the brain and CD68^+^ microglial burdens. Here, an interaction term between the brain region and DNAm score was included to test if any effects were region dependent. The same covariates and random effect as above were included. Models assessing blood-based signatures were additionally adjusted for the interval between their measurement and death. In each regression analysis, continuous variables were scaled to have a mean of zero and unit variance. We considered a statistical significance threshold of p<0.05. We additionally discuss how results change at a more conservative Bonferroni-corrected level of significance (p<0.05/41 = 0.001).

## 3. Results

### 3.1 Cohort demographics

*Post-mortem* details for each individual included in the study are presented in **Supplementary Table 1.** Summary statistics for each of the variables included in analyses is presented in **Table 1.** Age at death ranged from 77.6 to 82.9 years (mean=80.3, SD=1.56). Five of the 14 (36%) individuals were female.

**Table 1.**
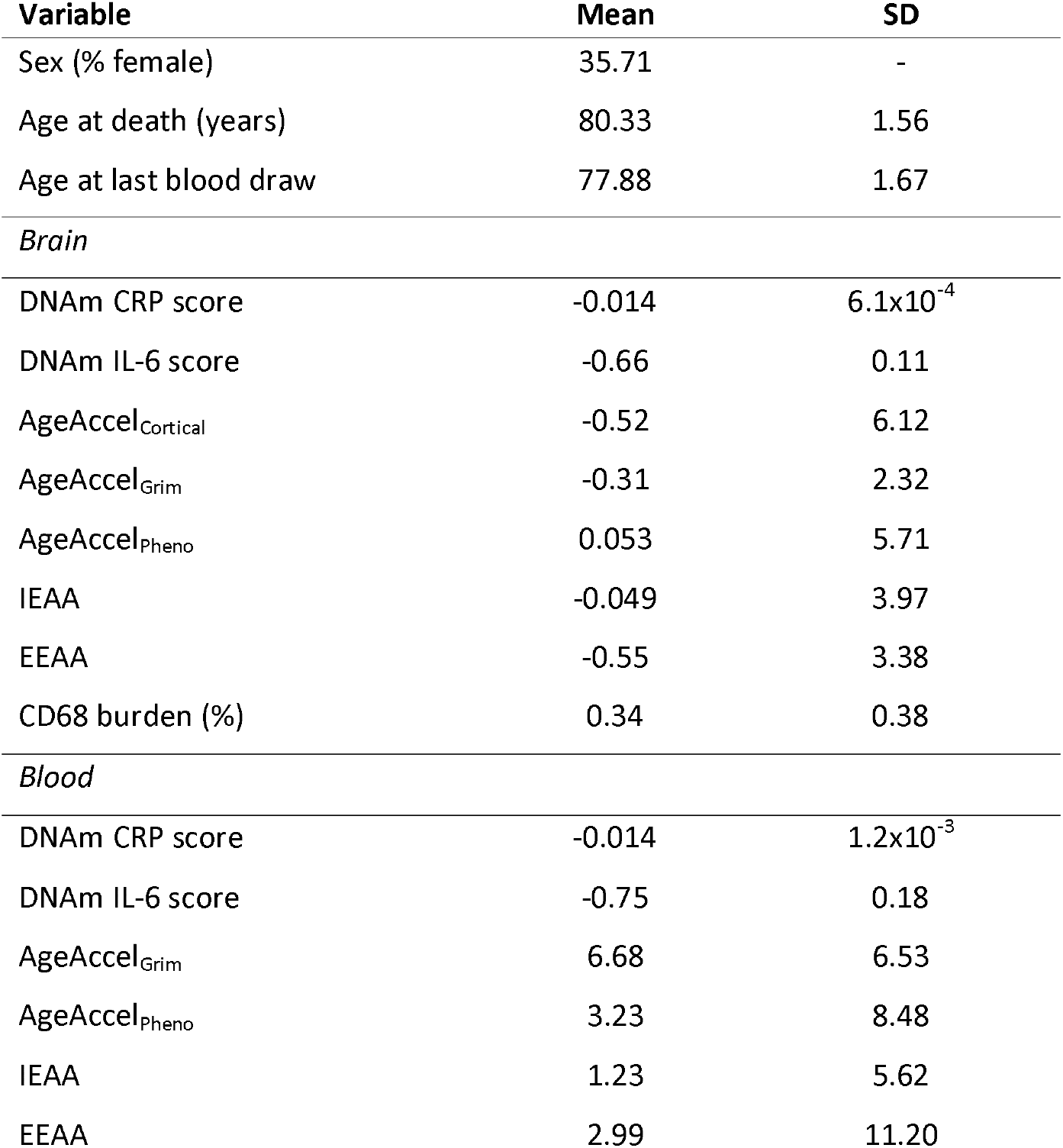
Summary of the variables assessed in the 14 Lothian Birth Cohort 1936 participants. The brain variables refer to the mean across all five regions. DNAm=DNA methylation; CRP= C-reactive protein; IL-6= interleukin-6; IEAA=intrinsic epigenetic age acceleration; EEAA= extrinsic epigenetic age acceleration; CD68=Cluster of Differentiation 68.

### 3.2 DNAm inflammation signatures

The Spearman correlation between the last blood DNAm CRP score and the mean brain DNAm CRP score was 0.06. This blood-brain correlation varied by region, ranging from −0.52 in BA17 to 0.46 in BA46 **(Supplementary Figure 1).**

A boxplot of the DNAm CRP score in the five brain regions is presented in **Figure 1.** No significant differences were identified in the analysis by region **(Supplementary Table 4),** indicating none of the assessed regions had a significantly different DNAm CRP score compared to BA17.

**Figure 1.**
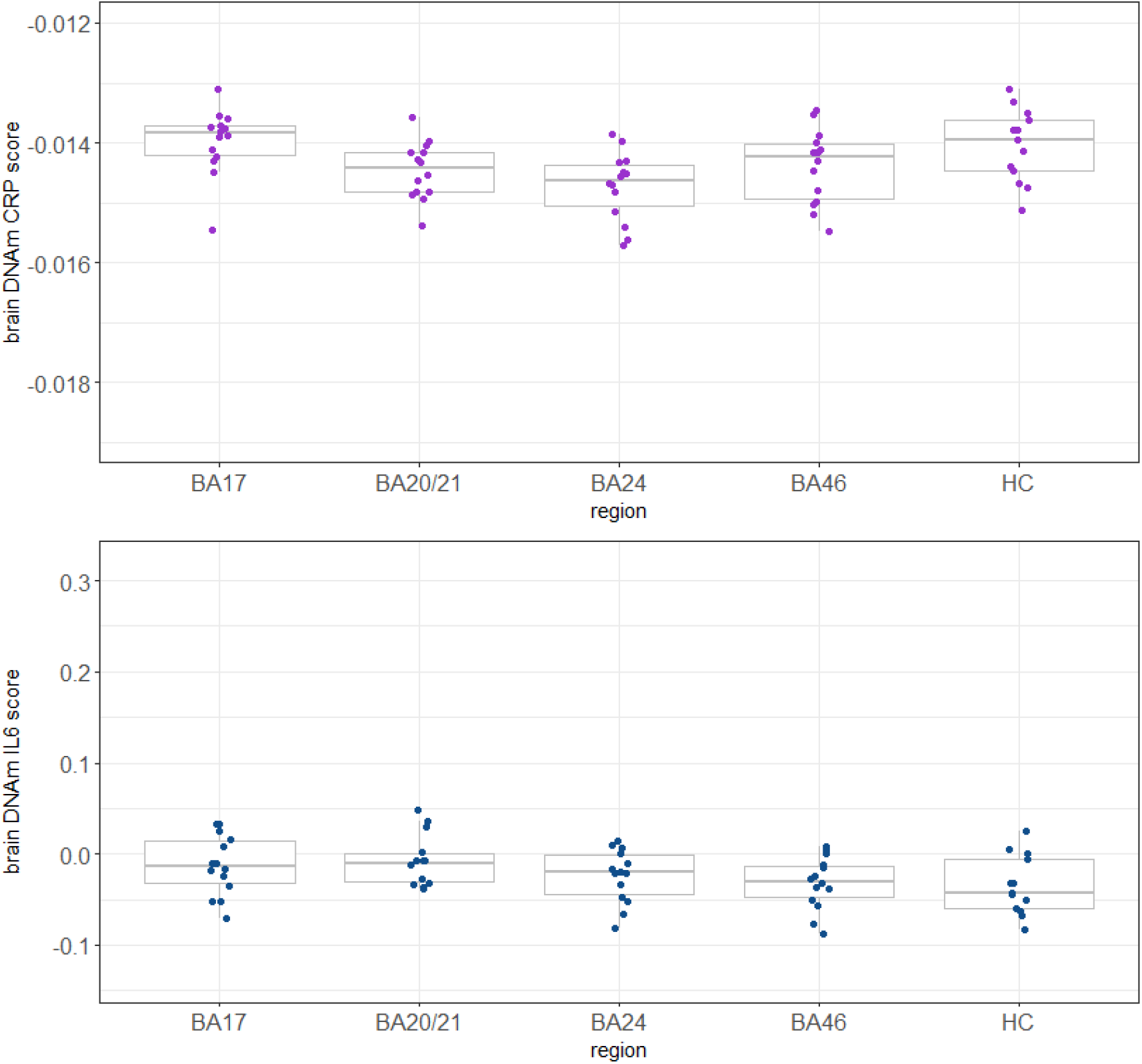
The DNAm CRP and IL-6 score in each of the five regions of the brain. BA=Brodmann area; HC=hippocampus; DNAm=DNA methylation; CRP=C-reactive protein; IL-6=interleukin-6.

The correlation between the last blood DNAm IL-6 score and the mean brain DNAm IL-6 score was 0.04, ranging from −0.12 in the hippocampus to 0.27 in BA46 **(Supplementary Figure 2).**

A boxplot of the DNAm IL-6 score in the five brain regions is presented in **Figure 1.** In the analysis by region, the DNAm IL-6 score was found to be significantly lower in BA24 (β=-0.86, SE=0.35, p=0.017), BA46 (β=-0.82, SE=0.30, p=0.009) and the hippocampus (β=-1.002, SE=0.32, p=0.003) compared to BA17 **(Supplementary Table 4).**

### 3.3 DNAm age acceleration

The correlations between the last blood DNAm age acceleration and the mean age acceleration in the brain were −0.04 for IEAA, 0.48 for EEAA, 0.39 for AgeAccel_Grim_, and 0.30 AgeAccel_pheno_. Correlation plots between the last blood DNAm age acceleration measure and the DNAm age acceleration in the brain split by region are presented in **Supplementary Figures 3-6.** The coefficients for AgeAccel_Grim_, AgeAccel_pheno_ and EEAA were all positive, ranging from 0.09 between AgeAccel_pheno_ in the blood and in BA46, to 0.78 between the last blood EEAA and EEAA in BA17. IEAA showed a negative correlation between the last blood measurement and the measure in BA20/21 (r=-0.27), BA24 (r=-0.14) and BA46 (r=-0.25) but a positive correlation in the hippocampus (r=0.30) and BA17 (r=0.49). For EEAA, some of the positive correlations appear largely driven by an individual with a high last blood measure (38.2) which corresponded with high measures in each of the brain regions **(Supplementary Figure 4).** This individual additionally had consistently high last blood measures in each of the other epigenetic age acceleration measures assessed (range: 6.6-25.4).

Boxplots of the five different epigenetic age acceleration measures in each of the five brain regions are presented in **Figure 2.** The hippocampus displayed the highest DNAm age acceleration compared to BA17 for each of the assessed measures except for AgeAccel_Grim_ which was highest in BA24 **(Supplementary Table 5;** AgeAccel_Cortical_: β=0.901, SE=0.19, p=2.6×10^-5^; AgeAccel_pheno_: β=1.14, SE=0.27, p=1×10^-4^; IEAA: β=0.83, SE= 0.34, p=0.02; EEAA: β=0.99, SE=0.24, p=1×10^-4^). The result for EEAA remained similar when the individual with consistently high measures across all regions was removed (β=1.22, SE=0.30, p=1.4×10^-4^).

**Figure 2.**
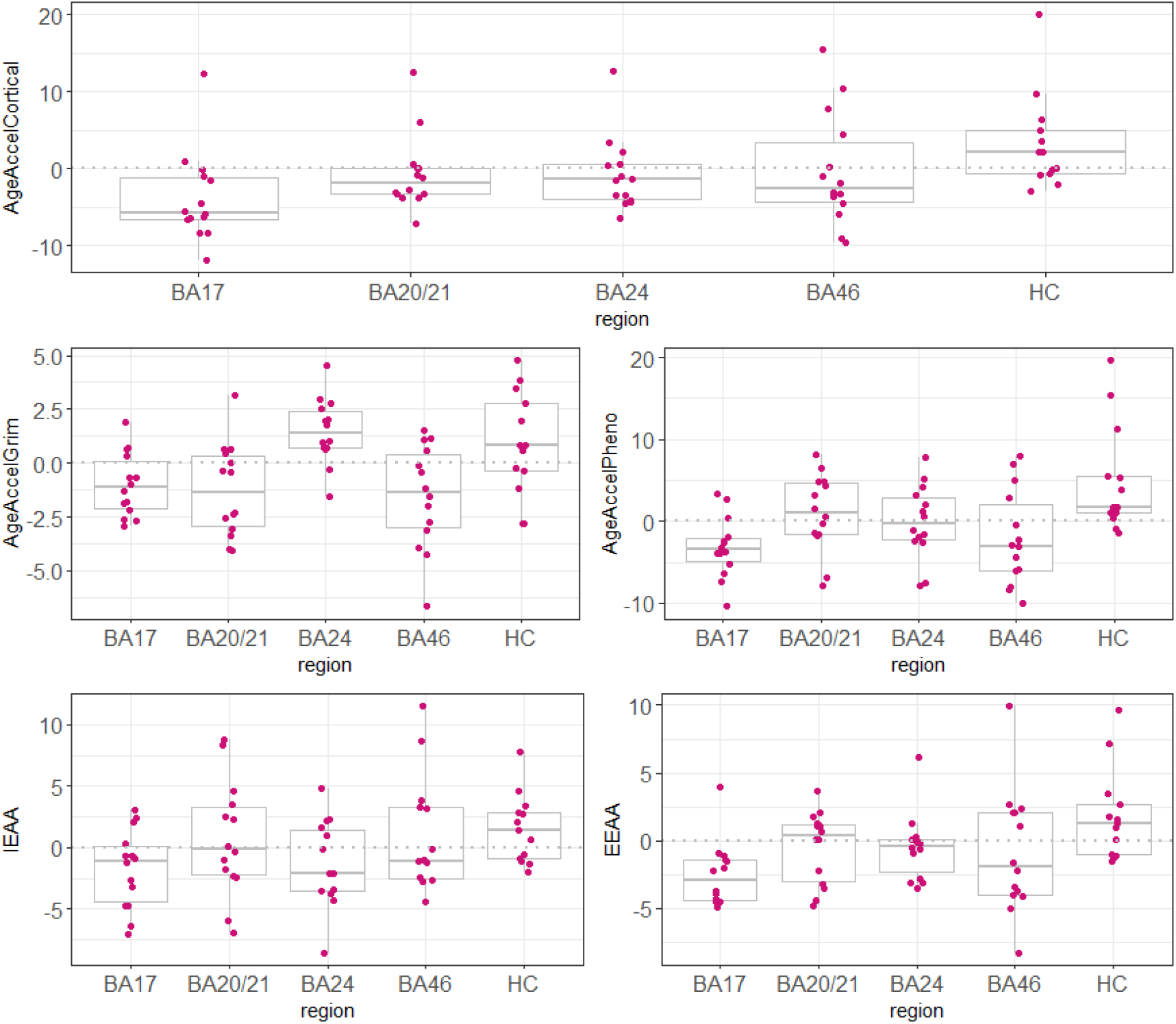
DNAm age acceleration measures across the five brain regions. The dashed grey lines represent where the mean difference is zero. IEAA=intrinsic epigenetic age acceleration; EEAA=extrinsic epigenetic age acceleration; BA=Brodmann area; HC=hippocampus.

### 3.4 Microglial burdens

A boxplot of the CD68^+^ microglial burdens in each of the five brain regions and a representative imaging of the staining is presented in **Figure 3.** The microglial burden was found to be significantly higher in the hippocampus compared to BA17 (β=1.32, SE=0.4, p=5×10^-4^), with the plot suggesting large variance in this region compared to the others.

**Figure 3.**
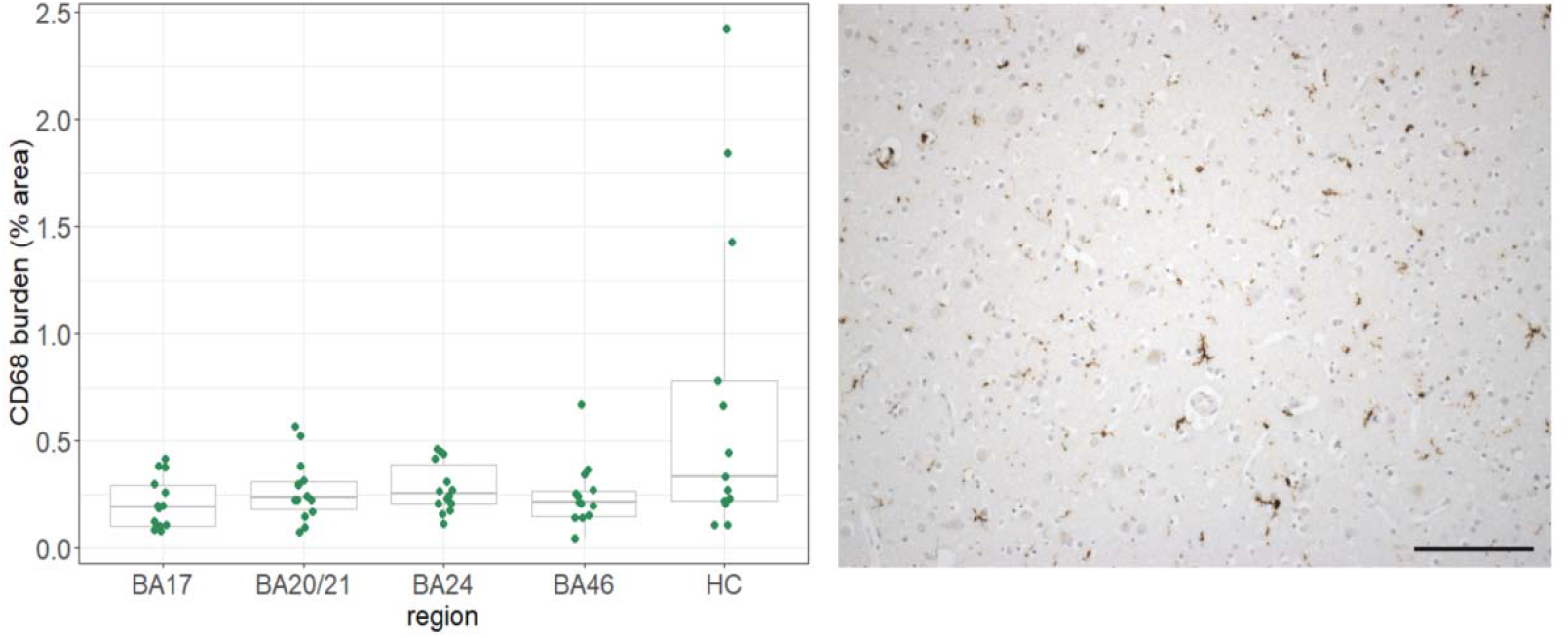
CD68^+^ microglial burdens over the five brain regions and representative staining. BA=Brodmann area; HC=hippocampus. Scale bar=150μm.

The associations between both the DNAm age acceleration variables and the DNAm inflammation signatures with microglial burdens are presented in **Supplementary Table 6.** Here, a higher mean AgeAccel_pheno_ in the brain associated with an increased microglial burden (β=0.40, SE=0.14, p=0.002). No other significant associations were identified (all p≥0.1) and there were no significant interactions found between any of the methylation scores and brain region.

## 4. Discussion

In this study, we took advantage of blood and *post-mortem* brain tissue available in 14 individuals in LBC1936 to investigate the relationship between peripheral and central inflammation- and age-related DNAm signatures and how they relate to neuroinflammatory processes. Due to the small sample size the results of this work are preliminary; however some potentially interesting patterns were identified. We found heterogeneous correlations between both the age acceleration, and inflammation-related, methylation signatures in the blood and the brain depending on the region assessed. Of the inflammatory signatures, the DNAm CRP score did not show significant variation across the brain regions, while the DNAm IL-6 score was found to be slightly lower in BA24, BA46 and hippocampus than in BA17. Other than for AgeAccel_Grim_, epigenetic age acceleration was found to be significantly higher in the hippocampus than in BA17. Reactive microglial burdens, identified through CD68 immunostaining, were additionally found to be higher in the hippocampus, consistent with previous findings in a smaller sample of the LBC1936 cohort (33). However, the only association identified between the DNAm signatures (age acceleration or inflammation proxies) and microglial load was a positive association with the mean brain-based DNAm AgeAccel_pheno_.

It is recognised that DNAm patterns at individual CpG sites in the blood and the brain are often disparate (34). We found that DNAm scores for CRP and IL-6 comprising multiple CpG sites displayed heterogeneous, region-specific correlations when comparing the blood- and brain-derived signatures. This suggests that blood DNAm patterns may proxy methylation in some areas of the brain better than others. Additionally, it cautions against the use of a single sample of post-mortem brain tissue as representative of the brain in aggregate, as it appears there is additional heterogeneity in methylation patterns even within the same tissue source. The DNAm age acceleration measures additionally displayed discrepant blood-brain correlations dependant on region. However, all the assessed measures showed positive blood-brain correlations in each region, to a greater or lesser degree, excepting IEAA. IEAA is based on the Horvath clock which is regarded as a pan-tissue model (11), whereas the other three peripheral measures were derived solely on blood DNAm data. Estimates from the Horvath clock have previously been found to be consistent across tissue types, making it surprising that IEAA showed the most inconsistent blood-brain correlation. A recent study has, however, suggested that the age prediction ability of the Horvath clock begins to deteriorate in older age (>60 years), possibly due to saturation of methylation levels at some loci (35). This may have impacted our results given both blood and brain tissue were gathered from 70 years onwards. The blood-brain correlations identified here suggest significant heterogeneity between the tissues, contingent on region; however, it should be noted that the mean interval between methylation assessed in the blood and in the brain was 2.5 years which reflects a period where methylation alterations are possible (36).

In the regional analyses of DNAm signatures in the brain, no real differences emerged in the assessment of the DNAm CRP score. On the other hand the DNAm IL-6 score seemed to be lower in BA24, BA46 and the hippocampus compared to BA17, possibly suggesting a disparity in the DNAm inflammation signatures across the brain. CRP itself does not typically cross the blood-brain barrier (BBB) although its pro-inflammatory effects may lead to an increased paracellular permeability of the BBB (37). Additionally, when using post-mortem blood tissue there is a possibility of blood contamination due to the lack of perfusion at *post-mortem.* Conversely, IL-6 can cross the BBB through the brain’s cirumventricular organs and is additionally expressed in the brain itself. However, the DNAm signatures of CRP and IL-6 were both created in blood and have not yet been validated in brain tissue. Work to assess other blood-calibrated predictors within in brain tissue is currently ongoing. It seems likely that brain tissue may exhibit different alterations in methylation in response to inflammation that were not captured by the two DNAm inflammatory marker proxies utilised here. In contrast to the inflammatory results, a higher DNAm age acceleration in the hippocampus was found for each of the assessed measures apart from AgeAccel_Grim_. This was true both for the cortex-specific clock as well as for the measures developed in the blood (AgeAccel_Pheno_ and EEAA) or in multiple tissues (IEAA). This consistency implies that the hippocampus may represent a region more susceptible to biological ageing than other areas of the neocortex. Age-related decline in hippocampal volume is well established (38) and it is one of the earliest, and most profoundly, affected regions in Alzheimer’s disease, suffering insidious synapse loss and neuronal cell death culminating in a substantial atrophy as the disease progresses (39). While none of the individuals included in this study had a diagnosis of Alzheimer’s disease prior to their death, the hippocampus can suffer substantial deterioration before clinical dementia becomes evident. The accelerated epigenetic ageing noted here is perhaps capturing the vulnerability of this region.

Equivalent to this finding, we identified a higher percentage burden of CD68^+^ microglia in the hippocampus compared to BA17. CD68 is a marker of phagocytic activity and is typically used to classify reactive microglia. Microglia are important in the maintenance of integrity and function within the central nervous system; however, aged microglia have been shown to be more responsive to pro-inflammatory stimuli compared to the naïve cell-type. This altered phenotype can lead to exaggerated neuro-inflammation in response to peripheral or central immune challenges which can precipitate neuro-toxicity, and thus, degeneration (40, 41). The only association identified between the DNAm signatures and microglial load was a positive association with the mean brain AgeAccel_Pheno_; however we did not find any significant interaction between the DNAm signatures and region. The DNAm PhenoAge clock was trained on a set of nine haematological and biochemical measures that were found to optimally predict an individual’s ‘phenotypic age’ including four immune cell profiles (lymphocyte percent, mean cell volume, red cell distribution width and white blood cell count) alongside CRP and albumin (10). Despite being developed on blood DNAm data, the predominantly inflammatory and immune composition of this clock may mean that AgeAccel_Pheno_ is better able to capture process associated with inflammation even outwith the blood. In this regard, it may have outperformed the DNAm CRP and IL-6 score due to the inclusion of a composite set of phenotypes, which may more accurately index systemic inflammation compared to a single inflammatory surrogate.

This study provides a rarely-available assessment of data from blood, alongside *post-mortem* brain tissue methylation profiles and histology from the same individuals. Alongside this, profiling DNAm in multiple regions of the brain allowed us to investigate the heterogeneity of methylation patterns within the same tissue type. This study is limited by the small number of individuals for which data was available, leading to a lack of statistical power and the potential for both type 1 and type 2 errors. We considered p<0.05 as the threshold for statistical significance in the analyses. However, the following associations fail to pass a strict Bonferroni-corrected threshold (p≤0.001): the differences of the DNAm IL-6 score across the brain regions, the IEAA measure being highest in the hippocampus compared to BA17, and the association of DNAm AgeAccel_Pheno_ with the CD68+ microglia burden. This, again, highlights that the results presented here should be taken as preliminary patterns until analyses can be repeated in larger sample sizes. In regards to the microglial burdening, we used only one antibody (CD68) which limited definitive identification of labelled cells as parenchymal microglia. CD68 stains the lysosomes of ostensibly reactive microglia; however, the antibody can additionally stain infiltrating macrophages. Capturing both the microglia and macrophage burden still provides a relevant read-out of the cellular inflammatory status; however, further characterisation of the microglial phenotype, including generating a reactive:total ratio would be desirable to glean a better understanding of their specific relationship to DNAm signatures. Further to this, the burden metric used to quantify microglia could reflect differences in sizes of the cells as well as in total numbers. An additional aspect to bear in mind when utilising *post-mortem* tissue in methylation studies is the stability of global DNAm following death and the biological implications of this (42, 43). We attempted to account for the potential impact of this by adjusting analyses for *post-mortem* intervals; however as *post-mortem* changes in DNAm are not yet well characterised it cannot be ruled out that this confounded results. Finally, *post-mortem* studies will always be retrospective in nature, rendering it impossible to discern causal or consequential events.

In summary, using a well-characterised cohort of 14 individuals, we identified divergent correlations between the blood and brain in DNAm inflammation-related and age acceleration measures depending on region assessed. The hippocampus was found to display the highest DNAm age acceleration in four out of five assessed measures, potentially reflecting its inherent susceptibility to biological ageing and pathological processes compared to other cortical regions. The hippocampus additionally showed the highest burden of reactive microglia. Whilst an accelerated DNAm PhenoAge associated with an elevated microglial load across the brain, no region-specific associations were identified. Our results provide some initial indications of the blood-brain relationships in DNAm patterns and how these relate to central processes; however further work is needed to verify these results in larger sample sizes and to investigate how these patterns associate with cognitive function and neurodegenerative disease.

## 5. Acknowledgements

The authors thank all LBC1936 study participants and research team members who have contributed, and continue to contribute, to ongoing studies. LBC1936 is supported by Age UK (Disconnected Mind program) and the Medical Research Council (MR/M01311/1). Blood methylation typing was supported by Centre for Cognitive Ageing and Cognitive Epidemiology (Pilot Fund award), Age UK, The Wellcome Trust Institutional Strategic Support Fund, The University of Edinburgh, and The University of Queensland. This work was in part conducted in the Centre for Cognitive Ageing and Cognitive Epidemiology, which is supported by the Medical Research Council and Biotechnology and Biological Sciences Research Council (MR/K026992/1) and which supports IJD.

AJS and RFH are Translational Neuroscience PhD students funded by Wellcome (203771/Z/16/Z to AJS; 108890/Z/15/Z to RFH). REM and DLM_c_C are supported by an Alzheimer’s Research UK major project grant (ARUK-PG2017B-10). TSJ is supported by the European Research Council (ERC) under the European Union’s Horizon 2020 research and innovation programme (Grant agreement No. 681181) and the UK Dementia Research Institute which receives its funding from DRI Ltd, funded by the UK Medical Research Council, Alzheimer’s Society, and Alzheimer’s Research UK. PMV is supported by the Australian National Health and Medical Research Council (1113400) and the Australian Research Council (FL180100072). SRC is supported by the Medical Research Council (MR/R024065/1) and the US National Institutes of Health (R01AG054628).

