## Supplementary Figures 1-6 for "A comparison of blood and brain-derived ageing and inflammation-related DNA methylation signatures and their association with microglial burdens"

**Supplementary Figure 1.** Spearman correlations between the last blood DNAm CRP score and the DNAm CRP score in each of the five brain regions assessed. The correlation coefficients and 95% confidence intervals are presented in the table.

The age at last blood measurement ranged from 75.5 to 80.2 years and the age at death ranged from 77.6 to 82.9 years.

BA=Brodmann area; DNAm=DNA methylation; CRP=C-reactive protein

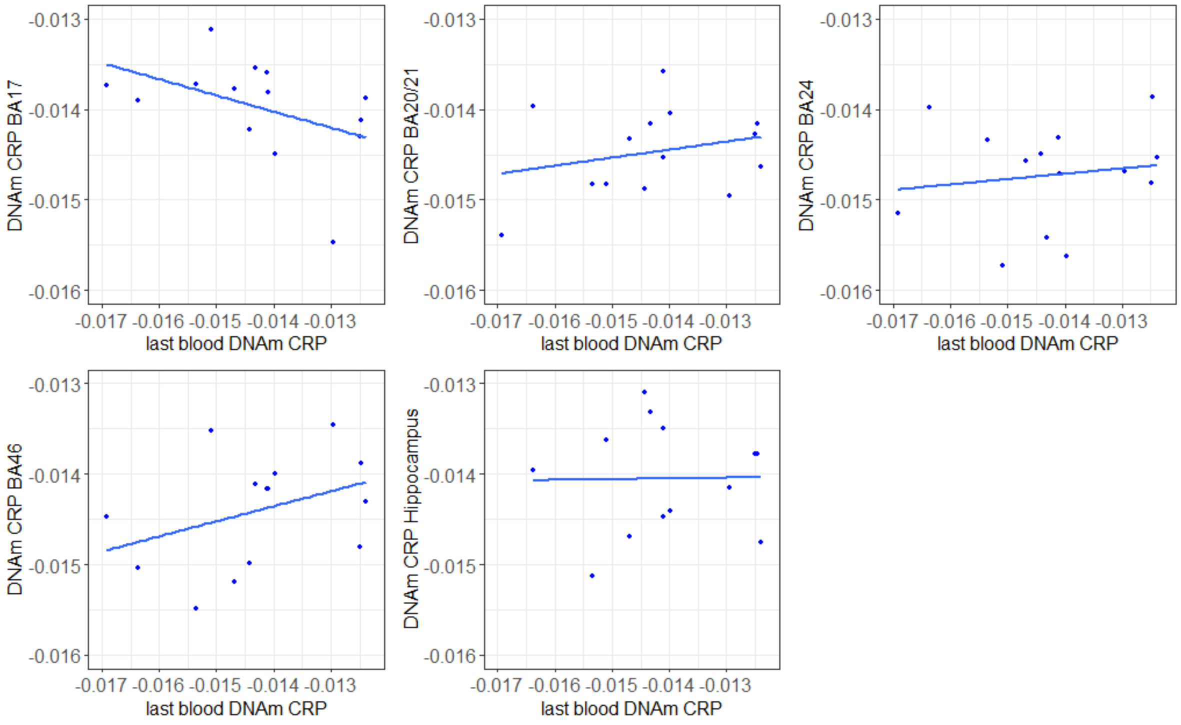


| Region | r | 95% confidence interval |
|---|---|---|
| BA17 | -0.52 | -0.79, 0.10 |
| BA20/21 | 0.20 | -0.31, 0.70 |
| BA24 | 0.07 | -0.42, 0.63 |
| BA46 | 0.46 | -0.20, 0.75 |
| Hippocampus | -0.07 | -0.54, 0.56 |

**Supplementary Figure 2.** Spearman correlations between the last blood DNAm IL-6 score and the DNAm IL-6 score in each of the five brain regions assessed. The correlation coefficients and 95% confidence intervals are presented in the table.

The age at last blood measurement ranged from 75.5 to 80.2 years and the age at death ranged from 77.6 to 82.9 years.

BA= Brodmann area; DNAm=DNA methylation; IL-6=interleukin-6

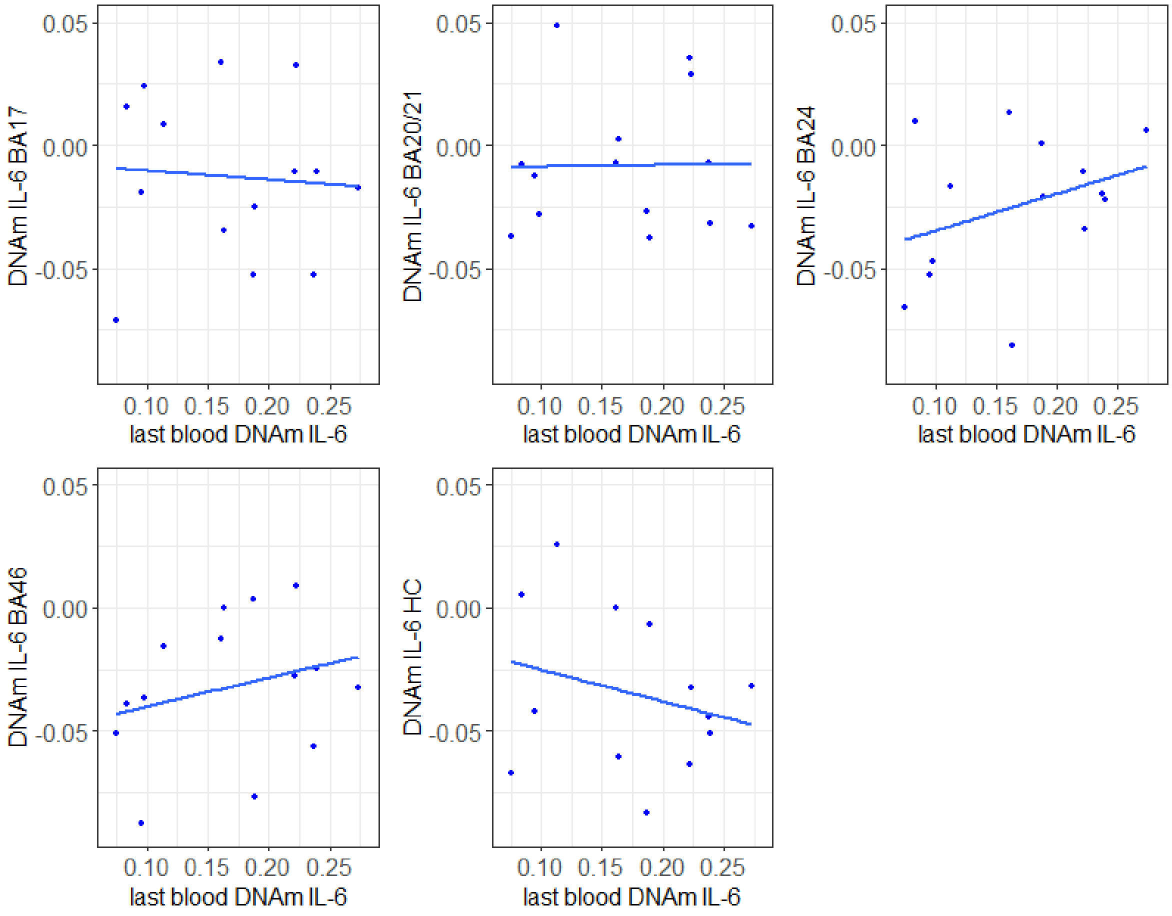


| Region | r | 95% confidence interval |
|---|---|---|
| BA17 | -0.01 | -0.58, 0.48 |
| BA20/21 | 0.02 | -0.52, 0.54 |
| BA24 | 0.22 | -0.23, 0.74 |
| BA46 | 0.27 | -0.31, 0.70 |
| Hippocampus | -0.12 | -0.71, 0.34 |

**Supplementary Figure 3.** Spearman correlations between the last blood IEAA and IEAA in each of the five brain regions assessed. The correlation coefficients and 95% confidence intervals are presented in the table.

The age at last blood measurement ranged from 75.5 to 80.2 years and the age at death ranged from 77.6 to 82.9 years.

BA= Brodmann area; IEAA=intrinsic epigenetic age acceleration

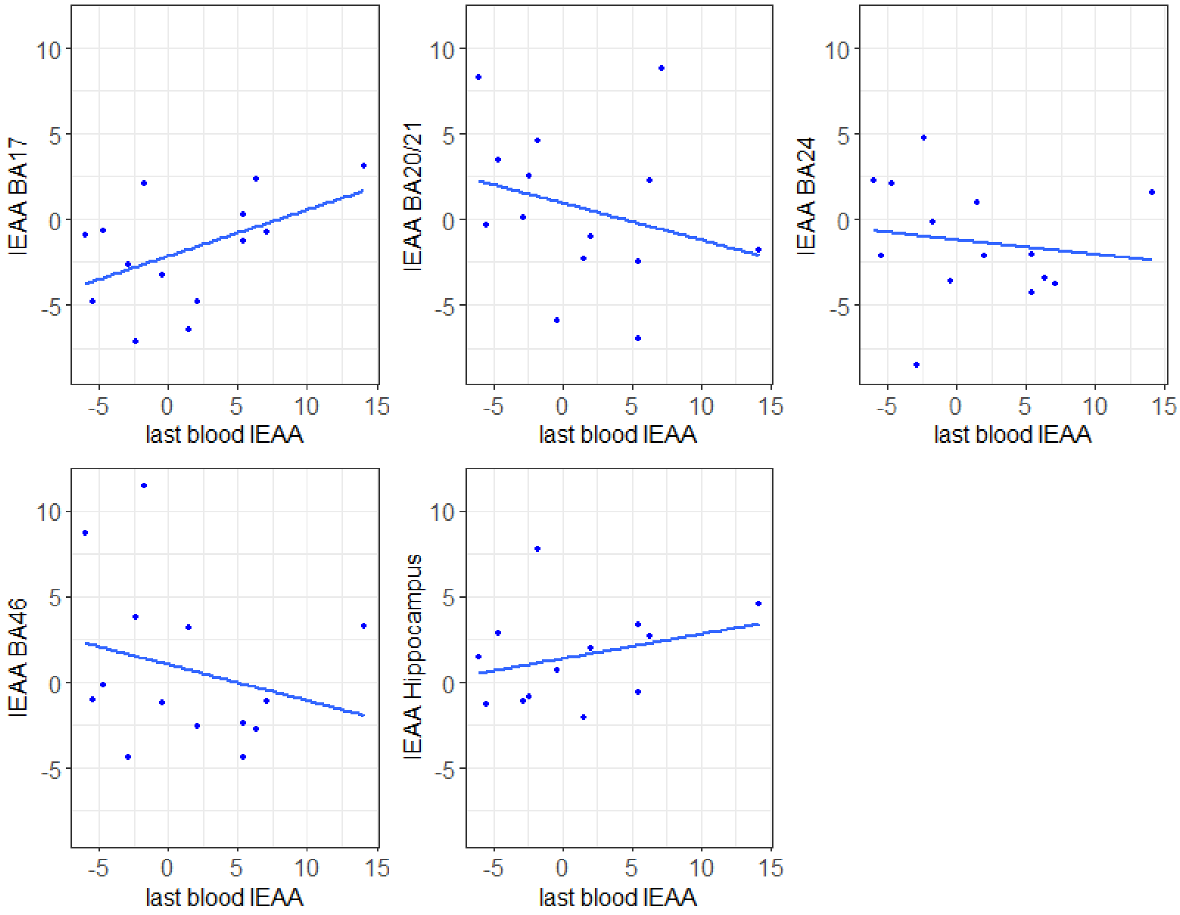


| Region | r | 95% confidence interval |
|---|---|---|
| BA17 | 0.49 | -0.058, 0.81 |
| BA20/21 | -0.27 | -0.69, 0.31 |
| BA24 | -0.14 | -0.63, 0.42 |
| BA46 | -0.25 | -0.69, 0.32 |
| Hippocampus | 0.30 | -0.30, 0.73 |

**Supplementary Figure 4.** Spearman correlations between the last blood EEAA and EEAA in each of the five brain regions assessed. The correlation coefficients and 95% confidence intervals are presented in the table.

The age at last blood measurement ranged from 75.5 to 80.2 years and the age at death ranged from 77.6 to 82.9 years.

BA= Brodmann area; EEAA=extrinsic epigenetic age acceleration

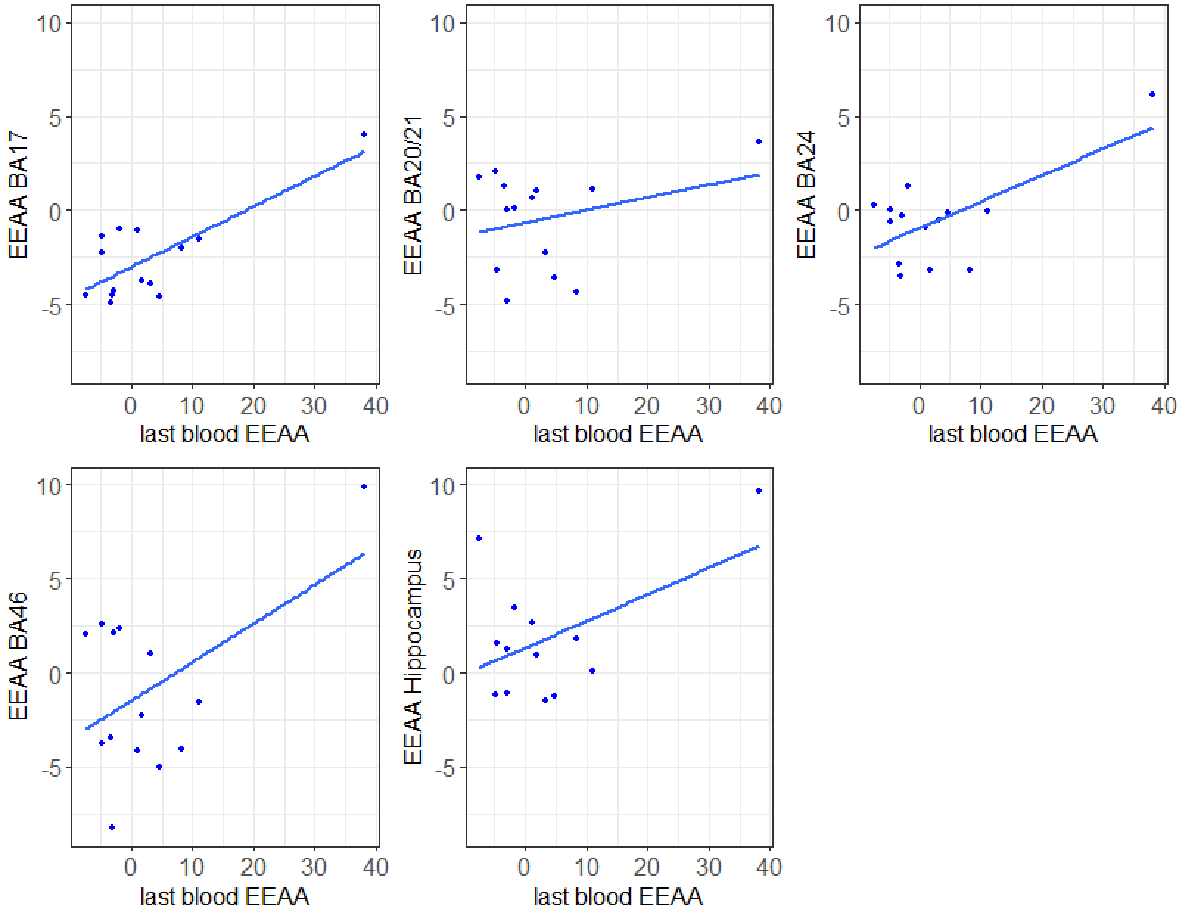


| Region | r | 95% confidence interval |
|---|---|---|
| BA17 | 0.78 | 0.42, 0.93 |
| BA20/21 | 0.29 | -0.29, 0.71 |
| BA24 | 0.66 | 0.20, 0.88 |
| BA46 | 0.52 | 0.019, 0.82 |
| Hippocampus | 0.50 | -0.069, 0.82 |

**Supplementary Figure 5.** Spearman correlations between the last blood AgeAccelGrim and AgeAccelGrim in each of the five brain regions assessed. The correlation coefficients and 95% confidence intervals are presented in the table.

The age at last blood measurement ranged from 75.5 to 80.2 years and the age at death ranged from 77.6 to 82.9 years.

BA=Brodmann area

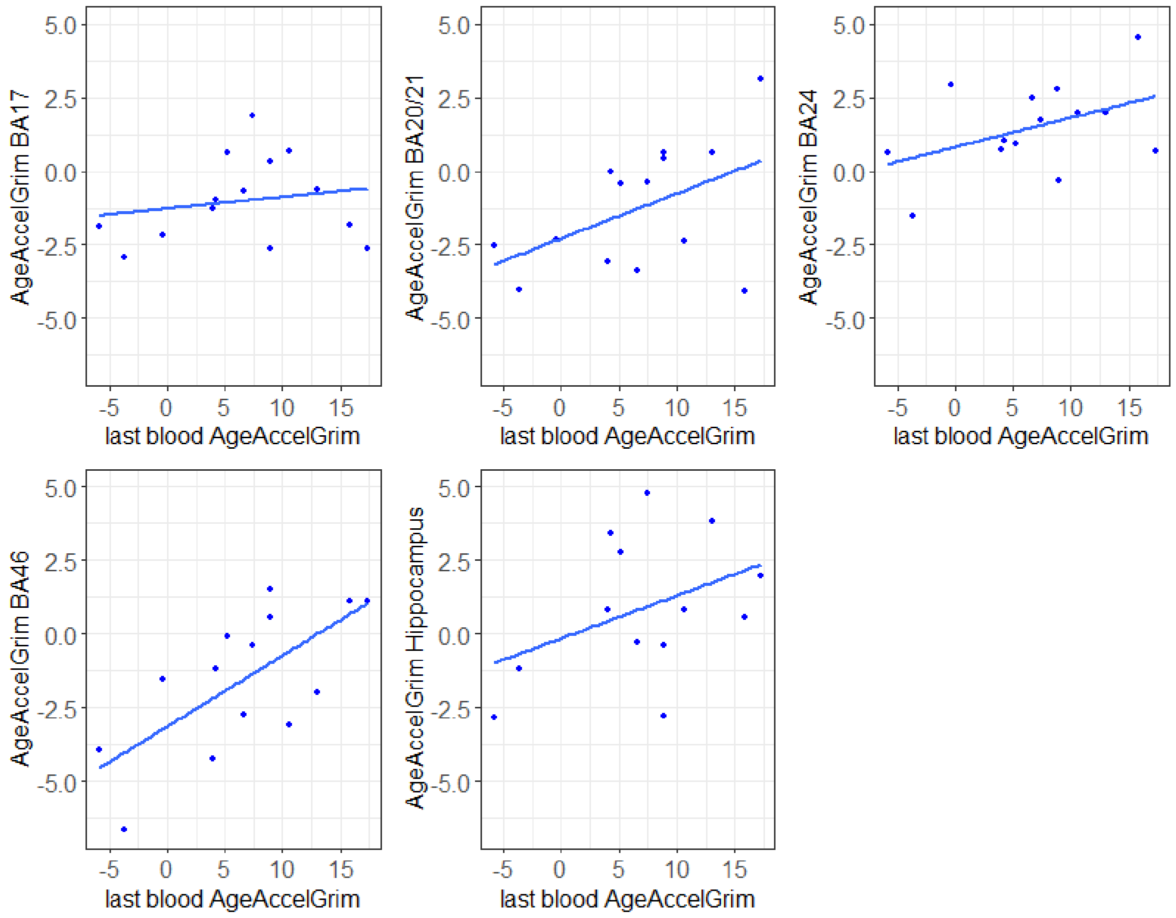


| Region | r | 95% confidence interval |
|---|---|---|
| BA17 | 0.18 | -0.39, 0.65 |
| BA20/21 | 0.48 | -0.069, 0.81 |
| BA24 | 0.44 | -0.11, 0.79 |
| BA46 | 0.68 | 0.23, 0.89 |
| Hippocampus | 0.40 | -0.19, 0.78 |

**Supplementary Figure 6.** Spearman correlations between the last blood AgeAccelPheno and AgeAccelPheno in each of the five brain regions assessed. The correlation coefficients and 95% confidence intervals are presented in the table.

The age at last blood measurement ranged from 75.5 to 80.2 years and the age at death ranged from 77.6 to 82.9 years.

BA=Brodmann area

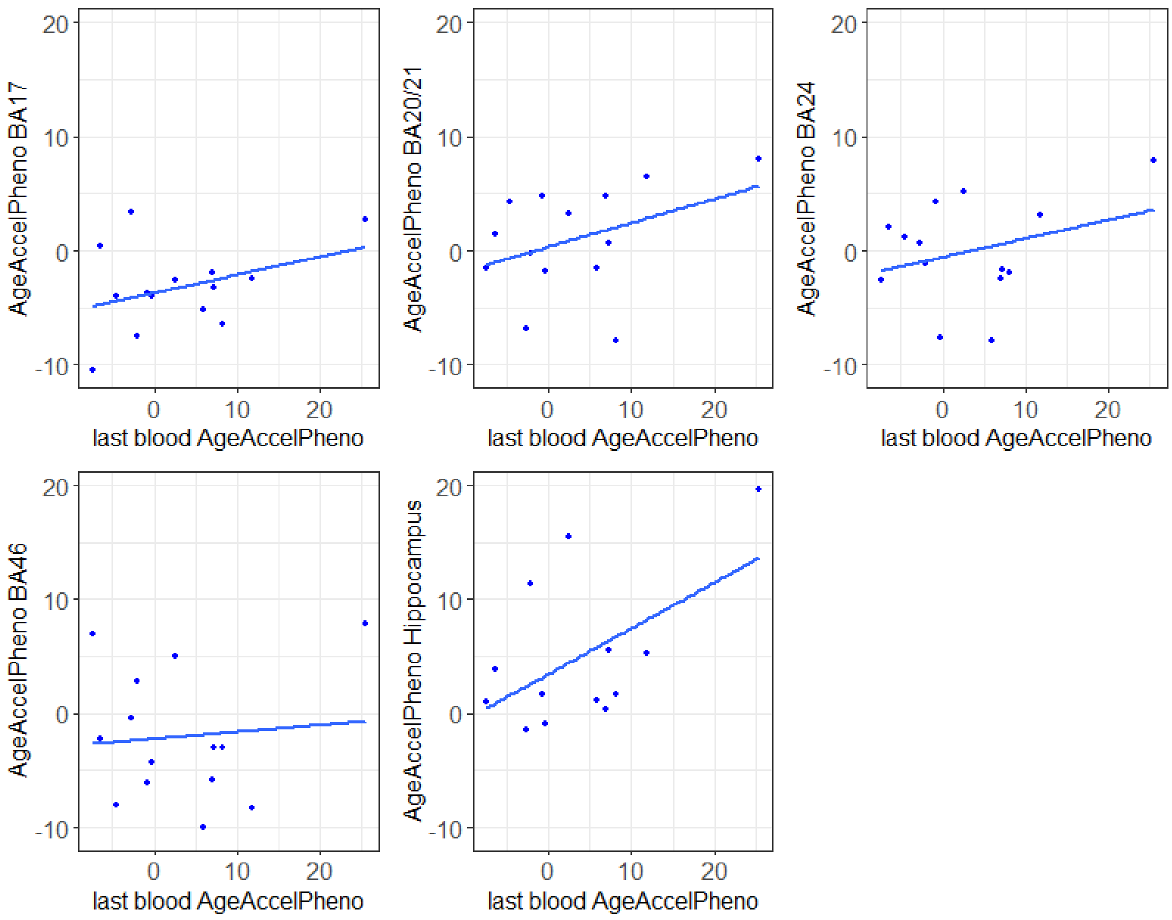


| Region | r | 95% confidence interval |
|---|---|---|
| BA17 | 0.37 | -0.20, 0.75 |
| BA20/21 | 0.39 | -0.18, 0.76 |
| BA24 | 0.32 | -0.26, 0.73 |
| BA46 | 0.09 | -0.46, 0.59 |
| Hippocampus | 0.54 | -0.021, 0.84 |
