## Supplementary Tables 1-6 for "A comparison of blood and brain-derived ageing and inflammation-related DNA methylation signatures and their association with microglial burdens"

**Supplementary Table 1.** *Post-mortem* details for the Lothian Birth Cohort 1936 participants.

CAA=cerebral amyloid angiopathy; COPD=chronic obstructive pulmonary disease; F=female; LVSD=left ventricular systolic dysfunction; M=male; PM=*post-mortem*; SVD=small vessel disease; WM=white matter.

| Brain bank number | Age at death | Sex | Interval between blood sample and death (years) | PM interval (hours) | Brain weight (g) | Brain pH | Cause of death | PM findings |
|---|---|---|---|---|---|---|---|---|
| BBN_19686 | 77 | F | 0.87 | 75 | 1320 | 6.5 | ·None specified | ·SVD<br>·Cerebral and cerebellar microinfarcts (lacunar)<br>·CAA<br>·Braak stage I |
| 001.26495 | 78 | M | 3.15 | 39 | 1290 | 6.2 | ·Amyloidosis<br>·Multiple Myeloma<br>·COPD | ·Braak stage I |
| 001.28402 | 79 | M | 2.29 | 49 | 1503 | 6.3 | ·Hospital acquired pneumonia<br>·Alcohol-related cirrhosis with ascites<br>·Severe LVSD<br>·COPD | ·Thal phase II<br>·Mild WM pathology<br>·Moderate non-amyloid SVD<br>·Hepatic encephalopathy<br>·Braak stage I |
| 001.28406 | 79 | M | 4.02 | 72 | 1437 | 6.1 | ·Suspected lung carcinoma<br>·Pneumonia | ·Thal phase II<br>·Moderate arteriolar CAA<br>·Braak stage II<br>·MildWM pathology<br>·Mild non-amyloid SVD |
| 001.28793 | 79 | F | 2.96 | 72 | 1219 | 5.9 | None specified | ·Thal phase I<br>·Moderate WM pathology<br>·Moderate non-amyloid SVD<br>·Braak stage II |

| | | | | | | | | |
|---|---|---|---|---|---|---|---|---|
| 001.28794 | 79 | F | 3.29 | 72 | 1289 | 5.8 | ·Metastatic breast cancer | ·Mild WM pathology<br>·Mild non-amyloid SVD<br>·Braak stage I |
| 001.28960 | 79 | M | 1.57 | 57 | 1301 | 6.1 | ·Chronic Lymphocytic Leukaemia<br>·Type 2 diabetes<br>·Myelofibrosis<br>·Idiopathic thrombocytopaenia purpura | ·Moderate WM pathology<br>·Moderate non-amyloid SVD |
| 001.29082 | 79 | F | 0.29 | 80 | 1339 | 5.9 | ·Metastatic carcinoma<br>·Squamous cell carcinoma lung | ·Thal amyloid phase V<br>·Mild arteriolar CAA<br>·Braak stage III<br>·Mild WM pathology<br>·Mild non-amyloid SVD |
| 001.29086 | 80 | F | 0.84 | 68 | 1468 | 6.2 | ·Intracerebral haemorrhage<br>·Hypertension | ·Cerebral haemorrhage<br>·Severe WM pathology<br>·Severe non-amyloid SVD |
| 001.31495 | 81 | M | 2.07 | 38 | 1318 | 5.8 | ·Stage IV Lung cancer<br>·Type 2 diabetes<br>·Ischaemic heart disease | ·Alzheimer disease<br>·Thal phase IV<br>·Braak stage VI<br>·Mild non-amyloid SVD<br>·Mild arteriolar CAA<br>·Mild WM pathology |
| 001.32577 | 81 | M | 1.18 | 74 | 1313 | 6.1 | ·Metastatic lung cancer<br>·COPD | ·Thal phase III<br>·Braak stage II<br>·Mild WM pathology<br>·Mild non-amyloid SVD<br>·WM microinfarcts<br>·Mild arteriolar CAA |

| | | | | | | | | |
|---|---|---|---|---|---|---|---|---|
| 001.34131 | 82 | M | 2.61 | 95 | 1472 | 5.9 | ·Squamous cell carcinoma of the lung | ·Thal amyloid phase III<br>·Severe arteriolar CAA<br>·Braak stage IV<br>·Severe WM pathology<br>·Moderate non-amyloid SVD |
| 001.35181 | 82 | M | 2.92 | 49 | 1496 | 6.1 | ·Bronchopneumonia<br>·Malignant neoplasm of the lung<br>·Acute renal failure<br>·Vascular dementia<br>·Ischaemic heart disease | ·Thal phase II<br>·Braak stage II<br>·Mild arteriolar CAA<br>·Mild WM pathology<br>·Mild non-amyloid SVD |
| 001.35215 | 82 | M | 6.05 | 40 | 1246 | 6 | ·Bronchopneuomina<br>·Left ventricular systolic dysfunction<br>·Hypertension<br>·Parotid abscess<br>·Prostate cancer<br>·Osteoarthritis | ·Braak stage I<br>·Severe WM pathology<br>·Severe non-amyloid SVD |

**Supplementary Table 2.** The DNAm CRP score probes and their coefficients**.** The CpG highlighted in bold was not available on the EPIC array and was omitted from analyses.

| ***CpG*** | ***coef*** |
|---|---|
| **cg06126421** | -0.0052 |
| cg06690548 | -0.0048 |
| cg10636246 | -0.0069 |
| cg18181703 | -0.0053 |
| cg19821297 | -0.0051 |
| cg25325512 | -0.0031 |
| cg27023597 | -0.005 |

**Supplementary Table 3.** The DNAm IL-6 DNA methylation probes and their coefficients.

| ***CpG*** | ***coef*** |
|---|---|
| cg04583842 | 0.04656555 |
| cg03885055 | -0.12047365 |
| cg17412005 | -0.328612128 |
| cg14965639 | -0.184598578 |
| cg14230378 | 0.13503161 |
| cg20789595 | -0.27792001 |
| cg19584649 | 0.096742747 |
| cg25250132 | 0.403243387 |
| cg04468741 | 0.017749147 |
| cg23729763 | 0.274636429 |
| cg20059928 | -0.008648995 |
| cg04928129 | -0.688776111 |
| cg12929678 | 0.20117534 |

**Supplementary Table 4.** The differences in the DNAm CRP and IL-6 scores over the brain regions. BA17 was used as the reference. Associations with $P<0.05$ are highlighted in bold.

DNAm=DNA methylation; CRP=C-reactive protein; IL-6=interleukin-6; BA=Brodmann area.

| | **β** | **SE** | **P** |
|---|---|---|---|
| *DNAm CRP score* | | | |
| BA20/21 | -0.14 | 0.32 | 0.65 |
| BA24 | -0.42 | 0.33 | 0.21 |
| BA46 | -0.35 | 0.29 | 0.23 |
| Hippocampus | 0.34 | 0.31 | 0.27 |
| *DNAm IL-6 score* | | | |
| BA20/21 | -0.25 | 0.33 | 0.46 |
| BA24 | -0.86 | 0.35 | **0.017** |
| BA46 | -0.82 | 0.30 | **$9.6 \times 10^{-3}$** |
| Hippocampus | -1.0083 | 0.32 | **$2.7 \times 10^{-3}$** |

**Supplementary Table 5.** The differences in the DNAm age acceleration measures over the brain regions.

BA17 was used as the reference. Associations with P<0.05 are highlighted in bold.

IEAA=intrinsic epigenetic age acceleration; EEAA=extrinsic epigenetic age acceleration; BA= Brodmann area.

| | **β** | **SE** | **P** |
|---|---|---|---|
| *AgeAccel*$_{Cortical}$ | | | |
| BA20/21 | 0.21 | 0.20 | 0.29 |
| BA24 | 0.14 | 0.21 | 0.52 |
| BA46 | 0.44 | 0.18 | **0.022** |
| Hippocampus | 0.90 | 0.19 | **$2.6\times10^{-5}$** |
| *AgeAccel*$_{Grim}$ | | | |
| BA20/21 | -0.56 | 0.29 | 0.055 |
| BA24 | 0.52 | 0.29 | 0.085 |
| BA46 | -0.46 | 0.26 | 0.090 |
| Hippocampus | 0.46 | 0.28 | 0.10 |
| *AgeAccel*$_{Pheno}$ | | | |
| BA20/21 | 0.34 | 0.28 | 0.23 |
| BA24 | 0.065 | 0.29 | 0.83 |
| BA46 | $5.5\times10^{-3}$ | 0.26 | 0.98 |
| Hippocampus | 1.14 | 0.27 | **$1.1\times10^{-4}$** |
| *IEAA* | | | |
| BA20/21 | 0.60 | 0.35 | 0.091 |
| BA24 | 0.10 | 0.37 | 0.78 |
| BA46 | 0.63 | 0.32 | 0.057 |
| Hippocampus | 0.83 | 0.34 | **0.018** |
| *EEAA* | | | |
| BA20/21 | 0.25 | 0.25 | 0.32 |
| BA24 | 0.15 | 0.26 | 0.57 |
| BA46 | 0.31 | 0.28 | 0.18 |
| Hippocampus | 0.99 | 0.24 | **$1.3\times10^{-4}$** |

**Supplementary Table 6.** Associations between DNAm age acceleration and inflammation scores in the blood and the brain and CD68[+] microglial burdens.

The blood variables refer to the last measurement available prior to death. The brain variables refer to the mean across all five regions. Associations with P<0.05 are highlighted in bold.

CRP=C-reactive protein, DNAm: DNA methylation; IEAA=intrinsic epigenetic age acceleration; EEAA=extrinsic epigenetic age acceleration; IL-6=interleukin-6

| | **β** | **SE** | **P** |
|---|---|---|---|
| *Brain* | | | |
| DNAm CRP score | 0.05 | 0.12 | 0.66 |
| DNAm IL-6 score | 0.02 | 0.13 | 0.89 |
| $AgeAccel_{Cortical}$ | 0.17 | 0.16 | 0.29 |
| $AgeAccel_{Grim}$ | 0.16 | 0.13 | 0.21 |
| $AgeAccel_{Pheno}$ | 0.40 | 0.12 | **0.002** |
| IEAA | 0.18 | 0.13 | 0.18 |
| EEAA | 0.06 | 0.14 | 0.68 |
| *Blood* | | | |
| DNAm CRP score | -0.28 | 0.18 | 0.16 |
| DNA IL-6 score | 0.32 | 0.17 | 0.10 |
| $AgeAccel_{Grim}$ | 0.25 | 0.26 | 0.36 |
| $AgeAccel_{Pheno}$ | -0.17 | 0.21 | 0.45 |
| IEAA | -0.21 | 0.26 | 0.43 |
| EEAA | -0.25 | 0.19 | 0.21 |
